# SPaCeD uses spatial point process distances to pair heavy and light chains of B cell receptors from spatial BCR-seq

**DOI:** 10.64898/2026.04.19.719512

**Authors:** Yimeng Liu, Céline M. Laumont, Shreena Kalaria, Puneet Velidi, Zhengxiao Wei, Brad H. Nelson, Farouk S. Nathoo

## Abstract

We introduce SPaCeD, a framework that infers B cell receptor (BCR) heavy–light chain pairs from spatial BCR-seq by integrating expression matrices with spatial point process distances. In ovarian and breast cancer datasets and simulations, SPaCeD improves pairing recall and stability compared with existing methodology (*Repair*), particularly for pairs with lower spatial expression. More broadly, SPaCeD provides a framework for representing spatial omics data as spatial statistical objects.

---

Tumor-infiltrating B cells and plasma cells (TIL-Bs) play a critical role in anti-tumor immunity, yet their antigen specificities remain largely unknown. Each B cell expresses a unique B-cell receptor (BCR) composed of paired heavy and light chains that jointly determine antigen binding. Reconstructing these pairs is essential for understanding clonal architecture and identifying tumor-reactive antibodies. While single-cell RNA sequencing can recover heavy and light chain BCR pairs, it lacks spatial resolution and therefore cannot reveal how receptor clones are distributed or interact within the tumor microenvironment. Spatial transcriptomics (Ståhl et al. 2016; Bergenstråhle et al. 2020; Andersson et al. 2020) enables *in situ* measurement of BCR gene expression but does not directly identify which heavy and light chains belong to the same clone, as spatial drift can shift heavy and light chains so that they are not exactly co-located.

To our knowledge, existing methods specifically developed for heavy-light chain pairing from spatial transcriptomics data do not use spatial distance information. For example, Repair, (Engblom et al. 2023), relies solely on expression matrices while ImmunoMatch, (Guo et al. 2026), leverages antibody sequence features to predict pairing compatibility; however, both approaches do not explicitly account for spatial organization and distances between heavy and light chain spatial point patterns on the tissue. Related formulations that are widely used in spatial omics include Tangram, (Biancalani et al. 2021), CytoSPACE, (Vahid et al. 2023), novoSpaRc, (Nitzan et al. 2019), SpaOTsc, (Cang and Nie 2020), and moscot, (Klein et al. 2025), though these approaches are not specifically developed for heavy/light chain pairing from spatial BCR-seq.

A broader issue is that many computational approaches for spatial omics have been developed primarily by extending methods for non-spatial molecular data, with spatial information incorporated only indirectly or through relatively simple notions of neighborhood, Euclidean distance or co-localization. Yet spatial statistics provides a mature framework for treating the observed locations themselves as informative stochastic objects. In spatial BCR-seq, the locations at which a receptor clone is observed naturally define a spatial point pattern, and the similarity between two candidate receptor chains can therefore be assessed not only through their expression profiles but also through the geometry of their spatial distributions. The heavy–light pairing problem provides a particularly clear example in which this distinction matters: two chains arising from the same clone need not be exactly co-located because of spatial drift and measurement variability, but their point patterns should nevertheless remain spatially similar.

SPaCeD is a statistical framework that integrates expression and spatial distance information to infer heavy–light BCR pairings from spatial BCR-sequencing data. SPaCeD formulates pairing as a combinatorial optimization problem solved via a linear assignment algorithm, (Kuhn 1955; Munkres 1957), using an objective function that jointly considers the Repair-derived mapping matrix and spatial distance between clonal point patterns. In spatial transcriptomic data, multiple B cells may contribute to each spatial location, and clonally related B cells often occupy nearby regions within tissue niches. As a result, paired heavy and light chains from the same clone tend to exhibit similar spatial expression patterns, (Laumont et al. 2022; Phad et al. 2022). A schematic overview of the framework is shown in Fig. 1a–f.

**Figure 1:**
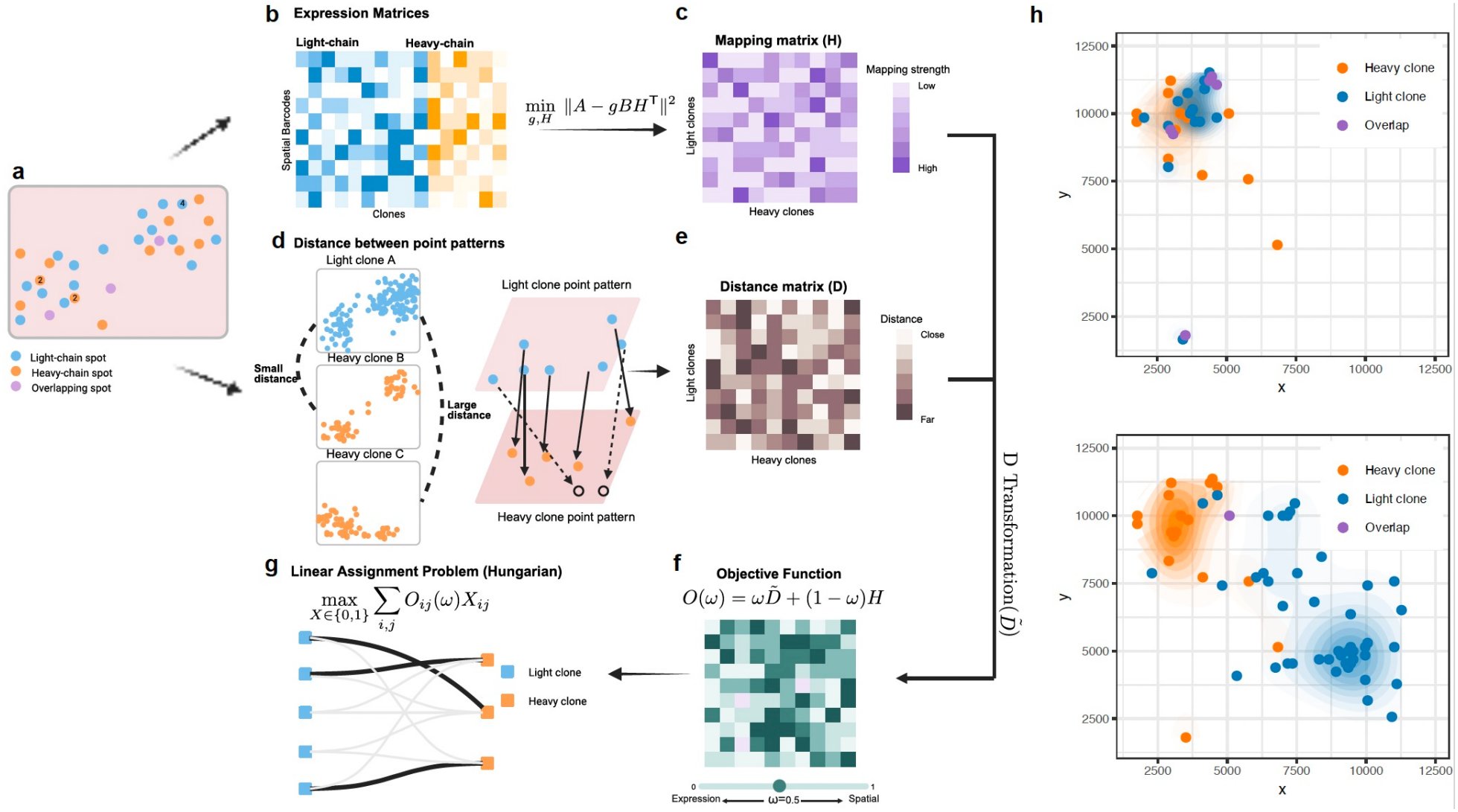
Workflow of SPaCeD and example overlay of light and heavy clones. **a**, Schematic illustration of spatial distributions of heavy- and light-chain expression spots. Blue and orange points denote light- and heavy-chain spots, respectively, with overlapping spots shown in purple. **b**, Example heavy-chain (*A*) and light-chain (*B*) expression matrices from spatial BCR-seq. **c**, An expression-based mapping matrix *H* is obtained by an optimization, quantifying relative expression contribution. **d**, Spatial information is incorporated by computing optimal-transport distances between heavy and light chain point patterns, yielding a distance matrix *D*. **e**, The distance matrix is transformed to 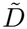. This transformation ensures that spatial distance is represented on the same scale as the expression-based mapping matrix, with larger values indicating stronger spatial agreement. **f**, The integrated objective matrix is formed as 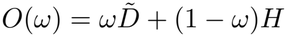, where *ω* ∈ [0, 1] modulates the influence of spatial versus expression information. **g**, The objective is supplied to a linear assignment solver (Hungarian algorithm) to infer heavy–light chain pairings. **h**, Spatial maps from the Engblom *et al*. breast cancer dataset showing a true pair (IGHGclone350– IGKCclone169; top) that SPaCeD identifies but Repair misses, characterized by closely aligned spatial distributions, and an incorrect pair (IGHGclone350–IGKCclone519; bottom) displaying discordant spatial patterns.

SPaCeD represents each heavy and light chain as a spatial point pattern and computes pairwise distances between these patterns using optimal transport. This yields a spatial similarity matrix that complements the expression-based mapping matrix obtained from Repair. The two sources of information are fused through a convex combination with a weighting parameter *ω* ∈ [0, 1], which controls the relative contribution of spatial versus expression information. Repair corresponds to *ω* = 0 (expression only), whereas *ω* = 1 relies exclusively on spatial point pattern distances. Instead of fixing *ω*, SPaCeD aggregates results across *ω* ∈ [0, 1], forming a multiscale solution of optimal assignments. For example, perfectly coincident pairs can be best identified by putting all of the weight on the expression information (*ω* = 0); whereas pairs that are not exactly coincident but spatially close will be better identified when more weight is placed on spatial distances. By construction, SPaCeD will include the Repair solution as a special case; however, it will further include pairs for which distance information can account for spatial drift.

To motivate the inclusion of spatial information, we examine two representative examples from the breast cancer dataset (Engblom et al. 2023) (Fig. 1h). A true heavy–light pair exhibits strongly overlapping and proximal spatial distributions, although the two point patterns are not perfectly collocated due to spatial drift. In contrast, an incorrect pair, despite having similar expression levels, shows clear spatial separation. Beyond the specific problem of BCR heavy and light chain reconstruction, our results demonstrate that explicitly modeling the spatial objects generated by spatial-omics experiments can provide information complementary to molecular abundance and can lead to substantial gains in downstream inference.

We conduct extensive simulation studies to evaluate the sensitivity and specificity of SPaCeD relative to Repair under varying degrees of noise and spatial drift. Simulations are generated using SRTsim, (Zhu et al. 2023), with spatial transcriptomic data from two ovarian cancer tissues (4A and 4C) and one breast cancer sample (Region C) as reference used to generate the simulated data. For each reference dataset, 50 replicates are produced under nine spatial settings, varying both the shift proportion *p* = 60%, 80%, 100%, defined as the fraction of heavy-chain points randomly selected for spatial drift, and a noise level *c ∈ {*5%, 15%, 25%*}*, defining the proportion of nearest on-tissue spots eligible as relocation destinations. In addition, we vary the clone-size threshold *k*, defined as the minimum number of spatial spots in which a clone must be observed to be retained for analysis. Unless otherwise stated, results reported in the main text correspond to *k* = 2, while results for *k* = 6 and *k* = 10 are provided in the Supplemental Material.

Across all datasets and noise levels, SPaCeD consistently outperforms Repair in pairing recall (Fig. 2b). As spatial perturbation increases, SPaCeD maintains relatively high recall, whereas the performance of Repair degrades sharply. Under the most extreme setting (*p* = 100%, *c* = 25%), SPaCeD achieves median recall values up to fourfold higher than those of Repair. These improvements are accompanied by an increase in the false-positive rate (FPR), particularly under greater spatial perturbation (Fig. 2c). Although this increase is noticeable in some simulation settings and should not be disregarded, the absolute FPR remains relatively low. This trade-off is observed across the simulation settings, with SPaCeD consistently achieving higher recall than Repair across all 27 conditions and clone-size thresholds *k* = 2, 6, 10 (Supplemental Figs. S1–S6).

**Figure 2:**
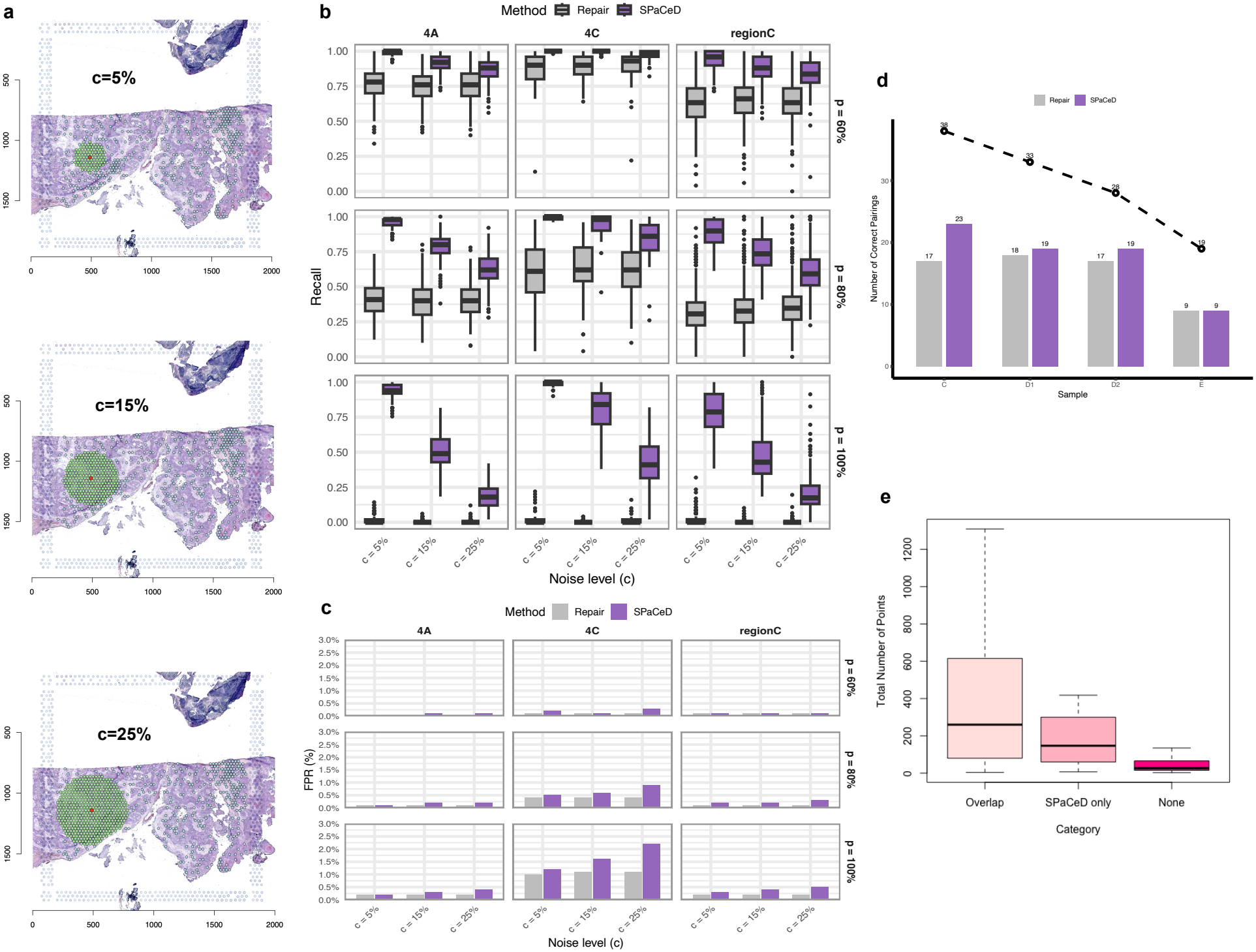
Evaluation of SPaCeD performance in simulated and real data. **a**, Schematic of the simulation design. Heavy-chain point patterns are perturbed by spatially shifting clone locations relative to their true light-chain partners. Three candidate relocation ranges are illustrated (noise level *c* = 5%, 15%, 25%). **b**, Recall of SPaCeD and Repair at clone-size threshold *k* = 2, stratified by shift proportion *p*. SPaCeD maintains high recall across all perturbation settings, whereas Repair shows increased sensitivity to larger values of *p*, with performance degrading as a greater fraction of heavy-chain points are spatially displaced. **c**, False-positive rates for SPaCeD and Repair under varying spatial noise. SPaCeD generally exhibits higher FPR than Repair, with the difference becoming more pronounced under greater spatial perturbation in some datasets. Nevertheless, FPR remains relatively low in absolute terms, indicating that the substantial improvements in recall are accompanied by a comparatively limited, though non-negligible, increase in false-positive pairings. **d**, Application to the Engblom *et al*. breast cancer dataset, showing the number of true heavy–light pairs identified in each tissue region. SPaCeD recovers at least as many ground-truth pairs as Repair in all four regions, with improvements observed in three regions. The dashed line indicates the total number of ground-truth pairs in each region. **e**, Distribution of the number of spatial spots supporting each ground-truth receptor pair, grouped by detection category: identified by both Repair and SPaCeD, identified exclusively by SPaCeD, or not identified by either method. SPaCeD uniquely recovers biologically plausible receptor pairs with moderate spatial signal, highlighting its improved sensitivity to low-abundance but spatially coherent clones.

We next apply SPaCeD to the breast cancer dataset of (Engblom et al. 2023), which includes four tissue sections (regions C, E, D1, and D2) with spatially resolved heavy- and light-chain expression. Ground truth heavy–light chain pairings derived from matched single-cell BCR sequencing are used to evaluate performance on these data (see, (Engblom et al. 2023), for details on ground truth pairs). SPaCeD recovers at least as many ground-truth heavy–light pairs as Repair across all four regions, with improvements observed in three regions. For example, in region C, SPaCeD identifies 23 ground-truth pairs compared with 17 for Repair, demonstrating improved recovery of true heavy–light chain pairs in real spatial BCR-seq data. These additional pairs tend to be spatially co-localized yet only moderately correlated in expression, suggesting that SPaCeD detects true matches that correlation-based methods fail to capture. To further characterize the improvements offered by SPaCeD over Repair, we examined the number of spatial spots contributing to each ground truth pair, and categorized pairs as detected by both methods, by SPaCeD only, or by neither.

As shown in Fig. 2e, pairs identified by both methods typically have larger spot counts, whereas pairs recovered by SPaCeD but not by Repair occupy an intermediate regime of moderate spatial abundance, indicating that SPaCeD can recover low-signal yet spatially coherent pairs that Repair fails to detect. Ground truth pairs undetected by both methods are generally associated with extremely sparse expression, limiting detectability for both algorithms.

As SPaCeD relies on distances computed within a single spatial region, its current implementation applies to contiguous tissue sections. This is not a major limitation in many applications, as most spatial BCR-sequencing studies analyze one section per sample. Nevertheless, the presence of consistent pairings across independently analyzed sections could serve as a prioritization criterion for downstream validation. Future work will extend SPaCeD to handle multi-section datasets, and work will also investigate the use of SPaCeD to infer T-cell receptor (TCR) *α*–*β* pairings, (Liu et al. 2022; Hudson and Sudmeier 2022; Sudmeier et al. 2022; Tu et al. 2019).

In summary, SPaCeD provides a simple yet powerful framework for integrating expression similarity and spatial organization to infer heavy and light receptor-chain pairs in spatial BCR-seq. In simulations, SPaCeD achieves up to fourfold higher pairing recall than Repair under severe spatial perturbation, while exhibiting a modest but non-negligible increase in false-positive rates.

## Methods

### Data and preprocessing

Spatial BCR-sequencing datasets were obtained from two ovarian cancer patients (4A and 4C) from the British Columbia Cancer Agency’s IROC study, and from the breast cancer dataset of (Engblom et al. 2023). The dataset contains multiple tissue regions; region C was used for simulation experiments, while all four regions were included in the real data analysis. Each dataset contained spatial barcode coordinates and spot-by-gene expression matrices for heavy- and light-chain transcripts. All analyses were performed in R (version 4.3.1).

### Expression-based receptor pairing with Repair

The Repair algorithm of (Engblom et al. 2023) serves as the baseline method for inferring heavy-light B-cell receptor (BCR) pairings from spatial BCR-seq data. Repair relies exclusively on gene expression similarity and does not incorporate spatial distance information. The method proceeds in two stages.

Let 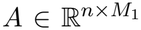 and 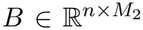 denote the spot-by-clone expression matrices for heavy and light chains, respectively. Repair estimates an unnormalized mapping matrix 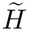and a global scaling factor 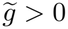 by solving the nonnegative regression problem

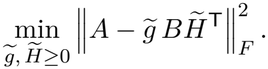

The matrix 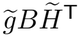 provides a low-rank reconstruction of heavy-chain expression using light-chain expression as predictors, and larger values 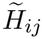 indicate stronger inferred expression-based association between heavy chain *i* and light chain *j*.

Because 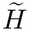 is unconstrained in scale, rows are normalized to obtain a row-stochastic mapping matrix

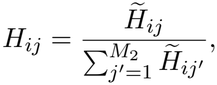

where 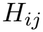reflects the relative expression-based contribution of light chain *j* to heavy chain *i*.

Given the normalized mapping matrix *H*, Repair formulates receptor pairing as a maximum bipartite matching problem. Let 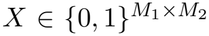 denote the assignment matrix, where *X*_*ij*_ = 1 indicates that heavy chain *i* is paired with light chain *j*. Repair solves

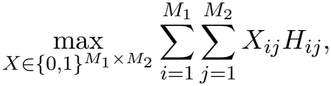

subject to the row-sum constraints. 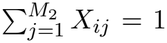 The optimal assignment is obtained using the Hungarian algorithm implemented in `solve_LSAP` (R package clue). Because Repair relies solely on *H*, pairing decisions depend entirely on expression similarity, and spatial distance information is not considered.

### Spatial distance between receptor clones

In spatial BCR-sequencing, receptor pairing is informed by two complementary signals: (i) expression similarity between heavy and light chains, and (ii) spatial proximity of their expression patterns across tissue. To quantify spatial similarity, each receptor clone is modelled as a spatial point pattern and distances between patterns are computed using optimal transport (OT).

Let *X* = *{x*_1_, …, *x*_*m*_*}* denote the spatial coordinates of spots expressing a heavy-chain clone and *Y* = *{y*_1_, …, *y*_*n*_*}* those expressing a light-chain clone, with *x*_*i*_, *y*_*j*_ ∈ R^2^. Each point pattern is treated as contributing one unit of mass, and we assume that this unit mass is distributed uniformly across the locations associated with each point pattern leading to mass distributions *p*_*i*_ = 1*/m* and *q*_*j*_ = 1*/n*. The OT distance was defined as the solution to

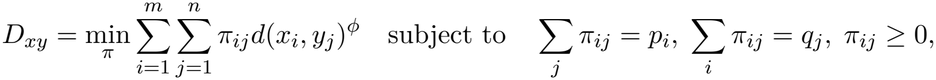

where *d*(*x*_*i*_, *y*_*j*_) denotes Euclidean distance and *ϕ* = 1 yields the Wasserstein-1 (Earth Mover’s) distance.

Because heavy and light chains often appear with different abundances and may exhibit partial overlap, we employed an unbalanced OT formulation implemented in the spatstat function pppdist. Unmatched mass was penalized using a cutoff distance *d*_cut_. Larger values of *d*_cut_ allow for long distance transport. For each heavy–light pair x, y, the resulting scalar OT cost was recorded as an entry *D*_*xy*_ of a spatial distance matrix 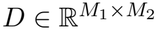. Smaller values of *D*_*xy*_ indicate closer spatial co-localization and greater biological plausibility of pairing.

### Integration of expression and spatial information in SPaCeD

Spatial distances between receptor clones were transformed into similarities via

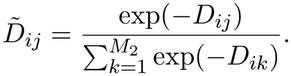

Expression-based similarity was represented by the normalized mapping matrix *H* estimated by Repair. The two sources of information were combined using a weighting parameter *ω* ∈ [0, 1] to define the SPaCeD matrix

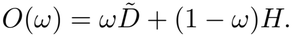

Here, *ω* controls the relative contribution of spatial distance versus expression-based evidence, with *ω* = 0 corresponding to Repair and larger values emphasizing spatial proximity to a greater degree.

For a fixed *ω*, heavy–light chain assignments are inferred by solving the linear assignment problem for *X*_*ij*_

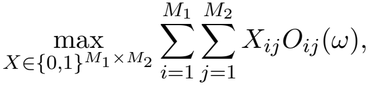

subject to the row-sum constraint 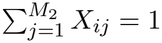. Optimization was performed using the Hungarian algorithm as implemented in solve_LSAP. This procedure was repeated for *ω* = 0, 0.1, …, 1. The final output of SPaCeD was defined as the union over *ω* solution, consisting of all heavy–light pairs that appear as optimal matches for at least one value of *ω*. Pair frequency across *ω* values and pairing multiplicity are used as indicators of robustness (Supplemental Table S1).

### Simulation design and performance evaluation

Synthetic spatial BCR-seq datasets were generated using SRTsim (Zhu et al. 2023), which uses real spatial transcriptomic datasets as references to simulate gene expression while preserving the underlying spatial structure. For each replicate, a light-chain expression matrix *L*^(*r*)^ was generated using SRTsim, and a binary one-to-one ground-truth pairing matrix *M* was randomly generated to define the true heavy–light chain pairs. Heavy-chain expression was then generated as

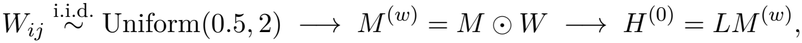

where *⊙* denotes element-wise multiplication and *H*^(0)^ represents the heavy-chain expression matrix before spatial displacement.

Spatial noise was introduced by randomly relocating a proportion *p ∈ {*60%, 80%, 100%*}* of heavy-chain expression locations. For each selected location, Euclidean distances to all on-tissue spots were calculated, and a destination was randomly selected from the nearest *c ∈ {*5%, 15%, 25%*}* of these spots. The relocated expression values were assigned to the corresponding destination spots in the heavy-chain expression matrix. For each combination of (*p, c, k*), 50 replicates were generated using three tissue templates: 4A and 4C (ovarian), and region C (breast).

Pairing performance was evaluated using recall and the false positive rate (FPR). Recall was defined as

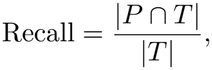

where *P* denotes the set of predicted pairs and *T* denotes the set of true pairs, corresponding to nonzero entries of *M* ^(*w*)^. The false positive rate was defined as

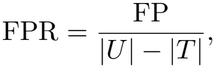

where FP denotes incorrectly predicted pairs and *U* is the set of all possible heavy–light chain combinations. Results were summarized using boxplots stratified by dataset and minimum clone size threshold *k*.

### Application to breast cancer spatial transcriptomics

SPaCeD and Repair are applied to the breast cancer spatial BCR-seq dataset from (Engblom et al. 2023) (regions C, E, D1, and D2). For each region, we compare the number of correctly recovered pairs and assessed spatial support for each inferred match. Pairs are categorized as detected by both methods, detected only by SPaCeD, or undetected. By construction, there cannot exist pairs detected by Repair that SPaCeD does not detect. SPaCeD consistently identifies additional spatially coherent receptor pairs with moderate spot abundance that Repair misses.

## Ethics and data provenance

This study involved secondary analysis of existing de-identified data obtained under appropriate data-access approvals/data-use agreements. No new participants were recruited and no new human samples were collected. Ethics approval and consent to participate for the original data collection were obtained by the original investigators, as described in the source studies.

## Software availability

SPaCeD is implemented in R and is publicly available at https://github.com/Floraliu7/Heavy-Light-Chain-Pairing. The repository provides the software used to infer heavy–light chain pairings from spatial BCR-seq expression data. Additional parameters and simulation settings are described in the Supplemental Material.

## Data access

The breast cancer spatial BCR-seq data analyzed in this study were previously published by Engblom et al. and are described in the Methods (European Genome-phenome Archive (EGA), accession number EGAD00001011061). Data and code for running the simulations and reproducing the ovarian and breast cancer simulation results are available at https://drive.google.com/file/d/10c2qNmqjafMVjtXonEQAzfHrp2xHGQCe/view?usp=sharing (ovarian) and https://drive.google.com/file/d/1oIUbRzEThzR4QpqXt4twsDvS4vz73UMV/view?usp=sharing (breast). SPaCeD source code is available at https://github.com/Floraliu7/Heavy-Light-Chain-Pairing.

## Competing interest statement

The authors declare no competing interests.

## Acknowledgments

Nathoo acknowledges funding from CIHR (PJT-186124), the Terry Fox Research Institute (TFRI1136 - 03) and NSERC (RGPIN-04044-2020). Nelson acknowledges the BC Cancer Foundation, CIHR (PJT-427647), and Terry Fox Research Institute (TFRI1060) for funding. Liu acknowledges funding from the Maud Menten Institute. We acknowledge the support of the Digital Research Alliance of Canada and the BC DRI Group for providing the research computing and storage resources used in this work. ChatGPT (OpenAI) was used to assist in identifying potentially relevant literature and copyediting (formatting, proof-reading, and minor wording suggestions for clarity). All retrieved references are independently verified by the authors, and all screening, interpretation, and citation decisions are made by the authors.

## Author contributions

Conceptualization: CML, BHN, FSN; Methodology: YL, FSN; Data analysis: YL; Data collection and preprocessing: CML, SNK, YL; Software: YL, ZW; Supervision: CML, BHN, FSN; Writing – original draft: YL, FSN; Writing – review and editing: CML, SNK, BHN, PV.

## Notes

https://github.com/Floraliu7/Heavy-Light-Chain-Pairing

### Competing Interest Statement

The authors have declared no competing interest.

### Summary of Updates

The title, abstract, and introduction have been revised to better emphasize the methodological contribution of SPaCeD and its broader relevance to spatial omics. The Methods section has been expanded to clarify the statistical framework, simulation design, and performance evaluation. Pairing accuracy has been renamed recall, and the false positive rate analysis has been expanded to better assess the trade-off between recall and specificity. The figures, supplementary materials, and discussion have been updated accordingly. The author list and author contributions have also been updated.

