## Supplementary Material for "SPaCeD uses spatial point process distances to pair heavy and light chains of B cell receptors from spatial BCR-seq"

### Contents

**Supplementary Tables** **2**

**Supplementary Figures** **4**

### List of Tables

S1 **Example of inferred heavy–light chain pairings.** Each row lists an inferred heavy–light chain pair. The *frequency* column denotes the number of $\omega$ values for which the pair is selected by SPaCeD, serving as a measure of pairing stability across different spatial–expression weightings. The *multiple heavy pair* column indicates whether a light chain is associated with multiple heavy chains, with a value of 1 indicating unique pairing to a single heavy chain. Together, these metrics provide a quantitative summary of pairing robustness and potential pairing ambiguity. . . . . . . . . . . . . . . . . . . . . . . . . . 2

S2 Specificity for Ovarian 4C simulation reorganized by spatial shift proportion $p$ and noise level $c$. . . . . . . . . . . . . . . . . . . . . . . . . . . . . . . . . . . . . . . . . 3

S3 Specificity for Ovarian 4A simulation reorganized by spatial shift proportion $p$ and noise level $c$. . . . . . . . . . . . . . . . . . . . . . . . . . . . . . . . . . . . . . . . . 3

S4 Specificity for Breast Region C simulation reorganized by spatial shift proportion $p$ and noise level $c$. . . . . . . . . . . . . . . . . . . . . . . . . . . . . . . . . . . . . . 3

### List of Figures

S1 **Recall across simulation settings ($k = 2$).** Boxplots of pairing recall for SPaCeD and Repair across three datasets (4A, 4C, and region C). Results are shown for different combinations of shift proportion $p$ (rows: $60\%, 80\%, 100\%$) and noise level $c$ (x-axis: $5\%, 15\%, 25\%$). Each box summarizes recall across 50 simulation replicates, with center lines indicating medians and boxes denoting interquartile ranges. SPaCeD maintains consistently high recall across all perturbation settings, whereas Repair performance degrades as spatial noise increases. . . . . . . . . . . . . . . . . . . . . . . . . . . 5

S2 **Recall ($k = 6$).** Same as Fig. S1, but for clone-size threshold $k = 6$. . . . . . . . . . . 6

S3 **Recall ($k = 10$).** Same as Fig. S1, but for clone-size threshold $k = 10$. . . . . . . . . . 7

S4 **False positive rate across simulation settings ($k = 2$).** False positive rate (1 − specificity) for SPaCeD and Repair across three datasets (4A, 4C, and region C). Results are shown for different combinations of shift proportion $p$ (rows: $60\%, 80\%, 100\%$) and noise level $c$ (x-axis: $5\%, 15\%, 25\%$). Each bar represents the mean false positive rate across 50 simulation replicates. Both methods maintain consistently low false positive rates across all perturbation settings. . . . . . . . . . . . . . . . . . . . . . . . . . . 8

S5 **False positive rate ($k = 6$).** Same as Fig. S4, but for clone-size threshold $k = 6$. . . . 9

S6 **False positive rate ($k = 10$).** Same as Fig. S4, but for clone-size threshold $k = 10$. . 10

### Supplementary Tables

Table S1: **Example of inferred heavy–light chain pairings.** Each row lists an inferred heavy–light chain pair. The *frequency* column denotes the number of $\omega$ values for which the pair is selected by SPaCeD, serving as a measure of pairing stability across different spatial–expression weightings. The *multiple heavy pair* column indicates whether a light chain is associated with multiple heavy chains, with a value of 1 indicating unique pairing to a single heavy chain. Together, these metrics provide a quantitative summary of pairing robustness and potential pairing ambiguity.

| **heavy_id** | **light_id** | **frequency** | **multiple heavy pair** |
|---|---|---|---|
| 125 | 1100 | 1 | 1 |
| 125 | 1434 | 10 | 1 |
| 126 | 1444 | 11 | 1 |
| 127 | 685 | 11 | 1 |
| 128 | 1557 | 11 | 1 |
| 129 | 1342 | 11 | 1 |
| 13 | 682 | 11 | 1 |
| 130 | 1458 | 11 | 1 |
| 131 | 1245 | 11 | 1 |
| 132 | 1270 | 11 | 1 |
| 133 | 1636 | 11 | 1 |
| 134 | 1512 | 11 | 1 |
| 135 | 346 | 11 | 1 |
| 136 | 786 | 11 | 1 |
| 137 | 108 | 11 | 1 |
| 138 | 1202 | 11 | 1 |
| 139 | 1088 | 11 | 1 |
| 14 | 22 | 11 | 1 |
| 140 | 717 | 11 | 1 |
| 141 | 493 | 11 | 1 |
| 142 | 764 | 11 | 1 |
| 143 | 266 | 11 | 1 |
| 144 | 1085 | 11 | 1 |
| 145 | 1157 | 9 | 1 |
| 145 | 1765 | 2 | 1 |

Table S2: Specificity for Ovarian 4C simulation reorganized by spatial shift proportion $p$ and noise level $c$.

| | | Cut 2 | | Cut 6 | | Cut 10 | |
|---|---|---|---|---|---|---|---|
| $p$ (%) | $c$ (%) | Repair | SPaCeD | Repair | SPaCeD | Repair | SPaCeD |
| **60** | 5 | 0.999 | 0.998 | 0.999 | 0.999 | 1.000 | 1.000 |
| | 15 | 0.999 | 0.999 | 0.999 | 0.999 | 1.000 | 0.999 |
| | 25 | 0.999 | 0.997 | 1.000 | 0.998 | 1.000 | 0.999 |
| **80** | 5 | 0.996 | 0.995 | 0.996 | 0.996 | 0.997 | 0.996 |
| | 15 | 0.996 | 0.994 | 0.996 | 0.995 | 0.997 | 0.996 |
| | 25 | 0.996 | 0.991 | 0.996 | 0.992 | 0.997 | 0.993 |
| **100** | 5 | 0.990 | 0.988 | 0.985 | 0.982 | 0.981 | 0.978 |
| | 15 | 0.989 | 0.984 | 0.984 | 0.976 | 0.979 | 0.970 |
| | 25 | 0.989 | 0.978 | 0.984 | 0.967 | 0.980 | 0.957 |

Table S3: Specificity for Ovarian 4A simulation reorganized by spatial shift proportion $p$ and noise level $c$.

| | | Cut 2 | | Cut 6 | | Cut 10 | |
|---|---|---|---|---|---|---|---|
| $p$ (%) | $c$ (%) | Repair | SPaCeD | Repair | SPaCeD | Repair | SPaCeD |
| **60** | 5 | 1.000 | 1.000 | 1.000 | 1.000 | 1.000 | 1.000 |
| | 15 | 1.000 | 0.999 | 1.000 | 1.000 | 1.000 | 1.000 |
| | 25 | 1.000 | 0.999 | 1.000 | 0.999 | 1.000 | 0.999 |
| **80** | 5 | 0.999 | 0.999 | 0.999 | 0.999 | 0.998 | 0.998 |
| | 15 | 0.999 | 0.998 | 0.999 | 0.998 | 0.998 | 0.997 |
| | 25 | 0.999 | 0.998 | 0.999 | 0.997 | 0.998 | 0.995 |
| **100** | 5 | 0.998 | 0.998 | 0.996 | 0.996 | 0.993 | 0.993 |
| | 15 | 0.998 | 0.997 | 0.996 | 0.994 | 0.993 | 0.989 |
| | 25 | 0.998 | 0.996 | 0.996 | 0.992 | 0.993 | 0.985 |

Table S4: Specificity for Breast Region C simulation reorganized by spatial shift proportion $p$ and noise level $c$.

| | | Cut 2 | | Cut 6 | | Cut 10 | |
|---|---|---|---|---|---|---|---|
| $p$(%) | $c$(%) | Repair | SPaCeD | Repair | SPaCeD | Repair | SPaCeD |
| **60** | 5 | 0.999 | 0.999 | 0.999 | 0.999 | 0.999 | 0.999 |
| | 15 | 0.999 | 0.999 | 0.999 | 0.999 | 0.999 | 0.999 |
| | 25 | 0.999 | 0.998 | 0.999 | 0.998 | 0.999 | 0.998 |
| **80** | 5 | 0.999 | 0.998 | 0.998 | 0.998 | 0.997 | 0.996 |
| | 15 | 0.999 | 0.998 | 0.998 | 0.997 | 0.997 | 0.996 |
| | 25 | 0.999 | 0.997 | 0.998 | 0.996 | 0.997 | 0.994 |
| **100** | 5 | 0.998 | 0.997 | 0.997 | 0.996 | 0.994 | 0.993 |
| | 15 | 0.998 | 0.996 | 0.996 | 0.993 | 0.994 | 0.989 |
| | 25 | 0.998 | 0.995 | 0.996 | 0.992 | 0.994 | 0.986 |

### Supplementary Figures

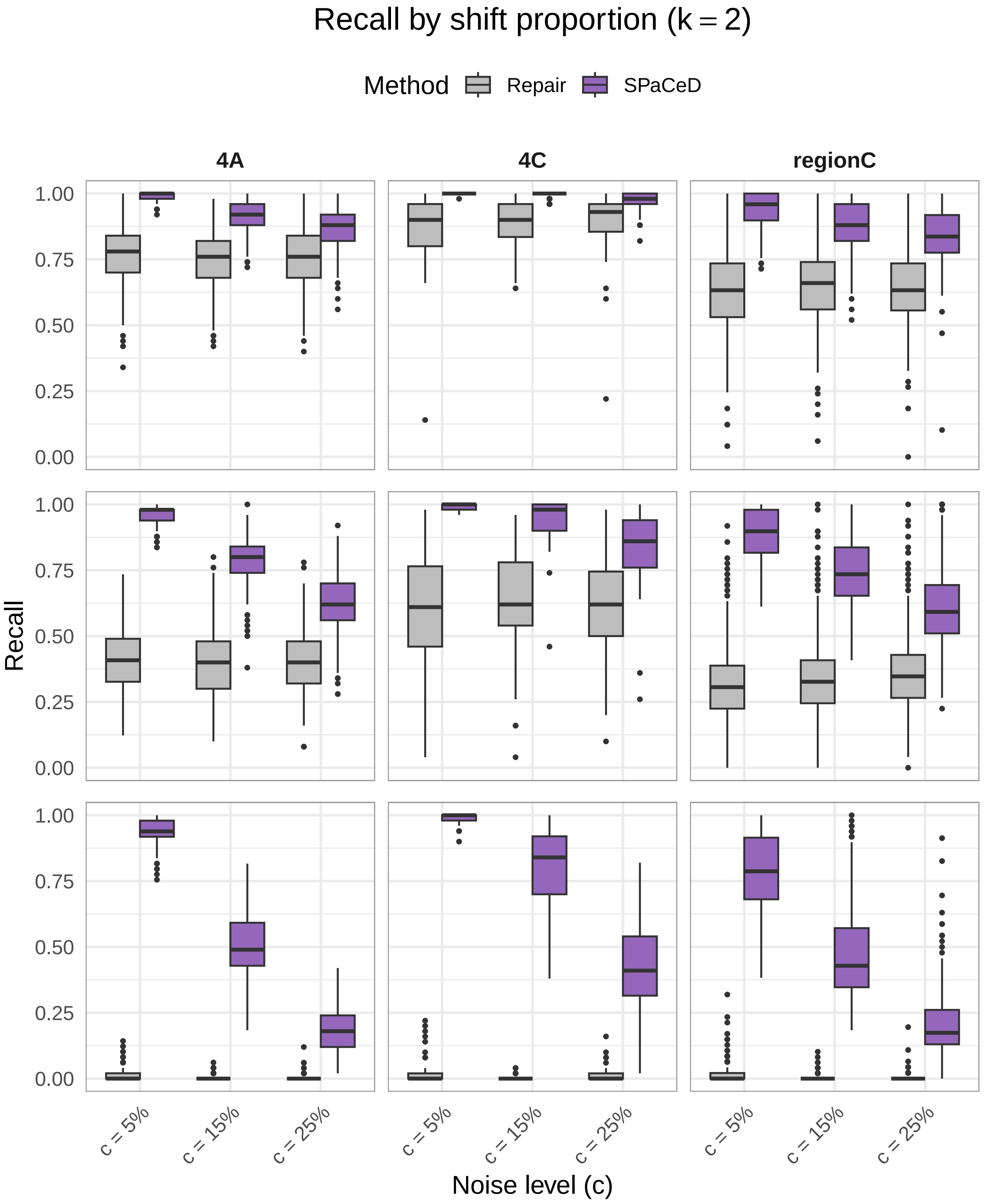


Figure S1: **Recall across simulation settings ($k = 2$).** Boxplots of pairing recall for SPaCeD and Repair across three datasets (4A, 4C, and region C). Results are shown for different combinations of shift proportion $p$ (rows: $60\%, 80\%, 100\%$) and noise level $c$ (x-axis: $5\%, 15\%, 25\%$). Each box summarizes recall across 50 simulation replicates, with center lines indicating medians and boxes denoting interquartile ranges. SPaCeD maintains consistently high recall across all perturbation settings, whereas Repair performance degrades as spatial noise increases.

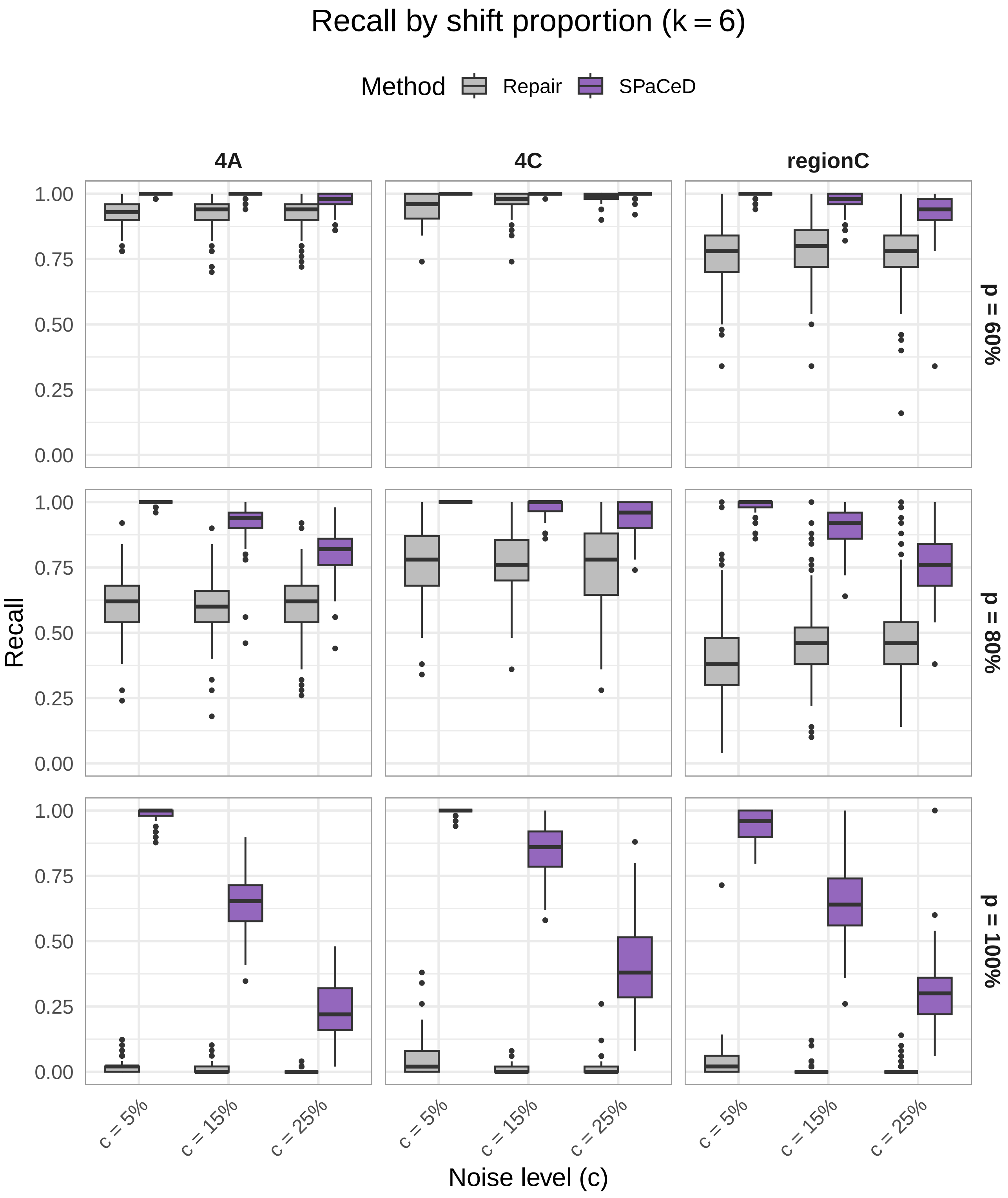


Figure S2: **Recall ($k = 6$).** Same as Fig. S1, but for clone-size threshold $k = 6$.

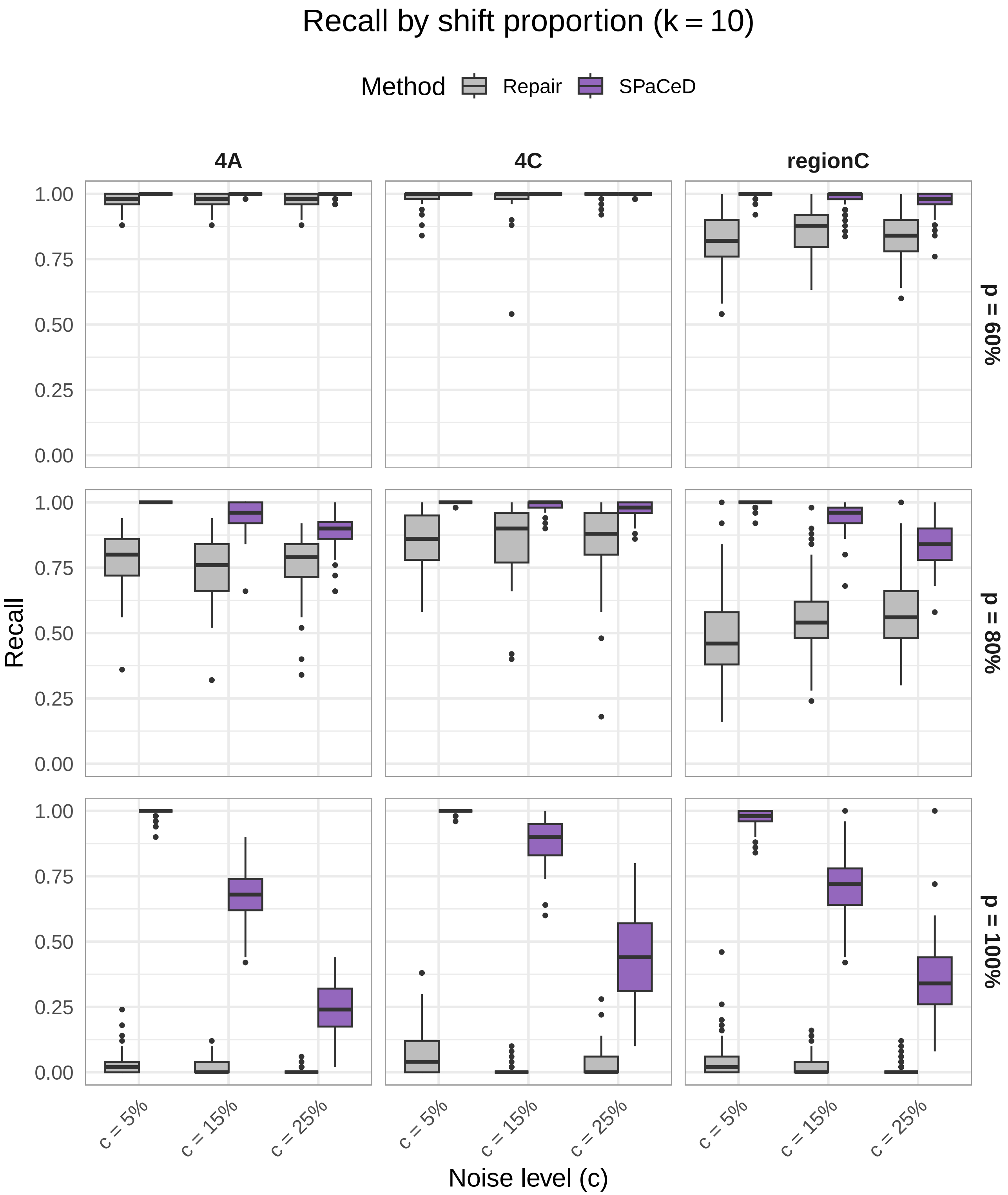


Figure S3: **Recall ($k = 10$).** Same as Fig. S1, but for clone-size threshold $k = 10$.

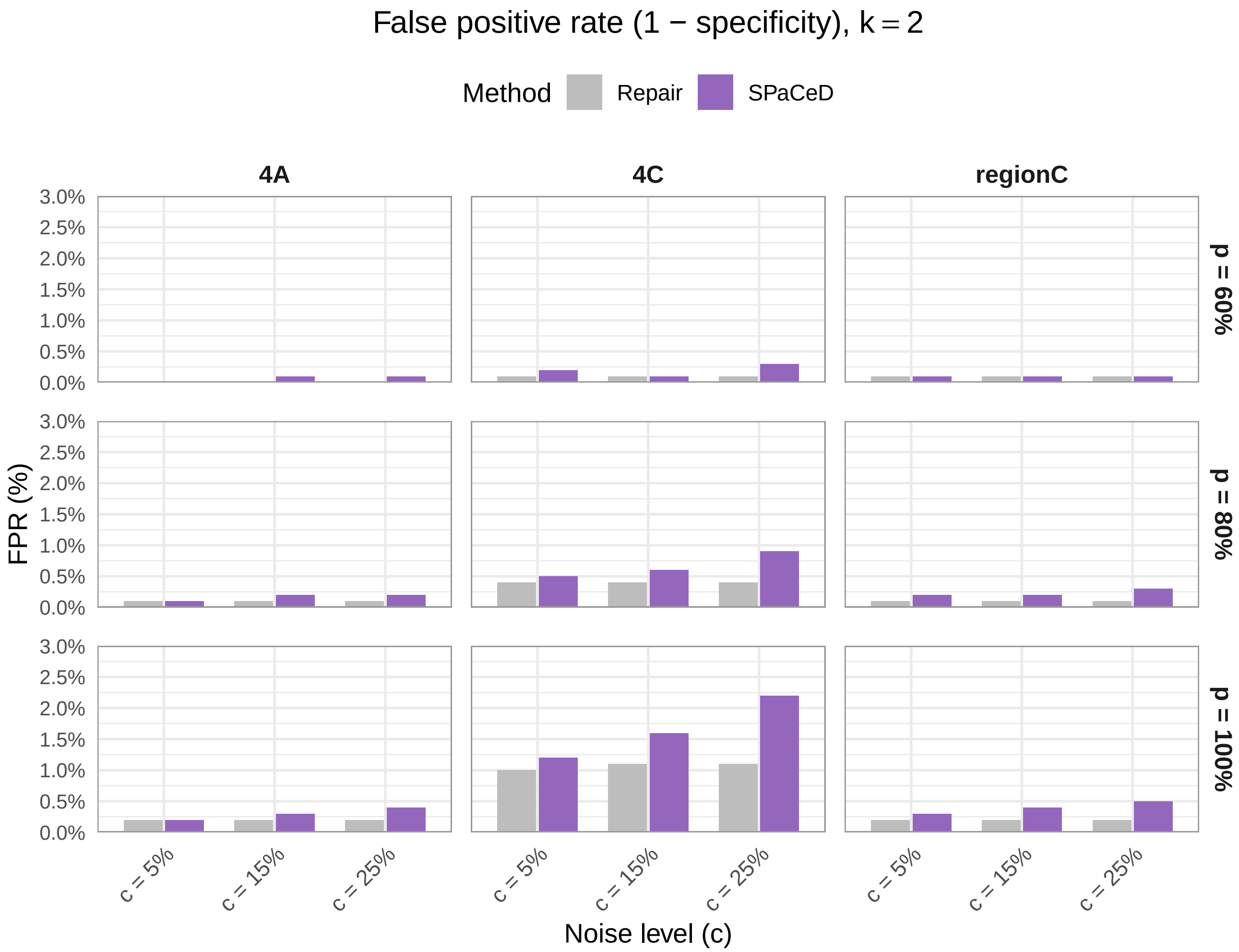


Figure S4: **False positive rate across simulation settings ($k = 2$).** False positive rate ($1 -$ specificity) for SPaCeD and Repair across three datasets (4A, 4C, and region C). Results are shown for different combinations of shift proportion $p$ (rows: $60\%, 80\%, 100\%$) and noise level $c$ (x-axis: $5\%, 15\%, 25\%$). Each bar represents the mean false positive rate across 50 simulation replicates. Both methods maintain consistently low false positive rates across all perturbation settings.

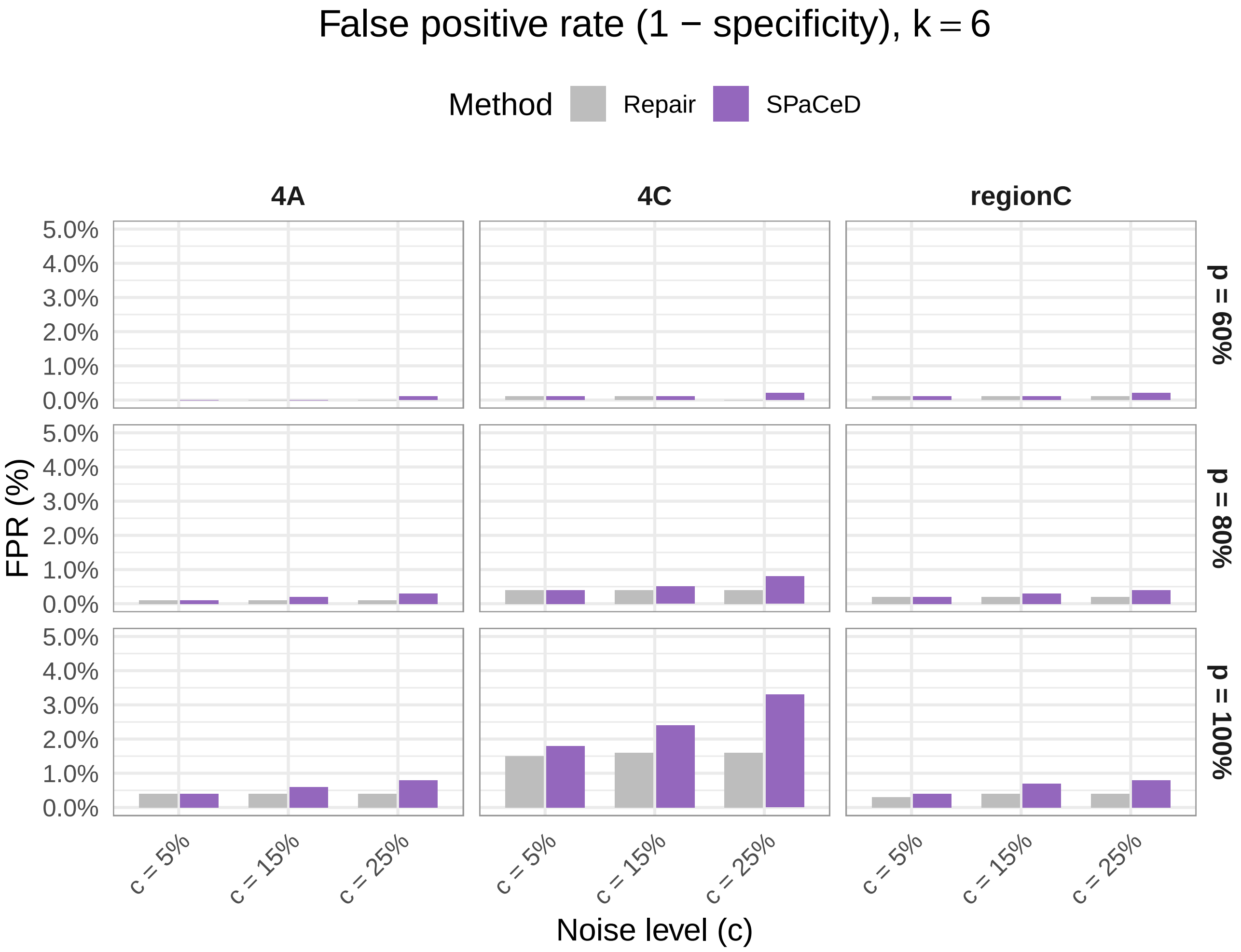


Figure S5: **False positive rate ($k = 6$).** Same as Fig. S4, but for clone-size threshold $k = 6$.

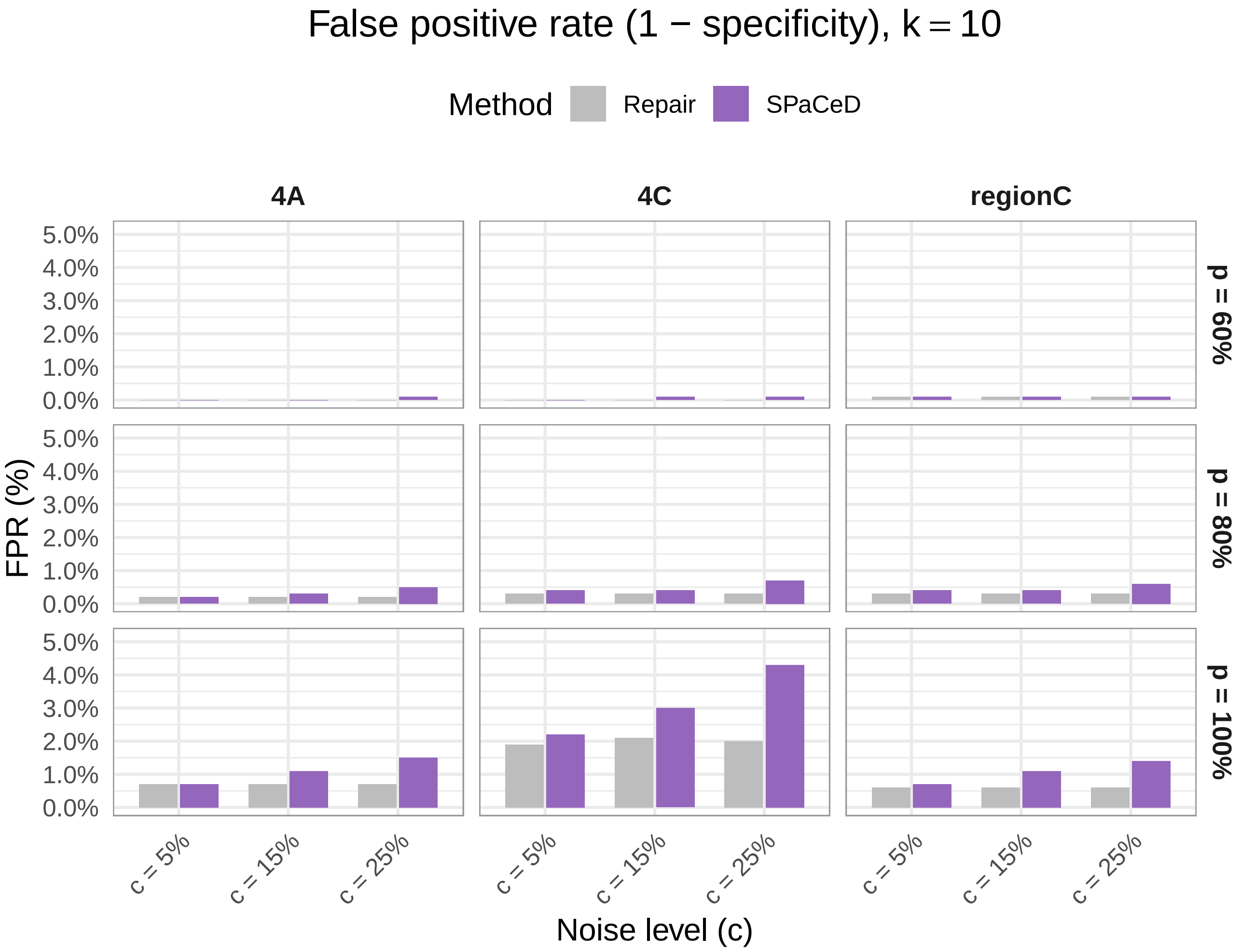


Figure S6: **False positive rate ($k = 10$).** Same as Fig. S4, but for clone-size threshold $k = 10$.

### Use of generative artificial intelligence

ChatGPT (OpenAI) was used to assist in identifying potentially relevant literature and copyediting (formatting, proof-reading, and minor wording suggestions for clarity). All retrieved references are independently verified by the authors, and all screening, interpretation, and citation decisions are made by the authors.
